## Supplementary material for "When two are better than one: Modeling the mechanisms of antibody mixtures": S1 Text

### Supporting information for When two are better than one: Modeling the mechanisms of antibody mixtures

### A Characterizing Antibodies Targeting the Receptor Tyrosine Kinase EGFR

#### A.1 Monoclonal Antibody Binding to a Receptor

In this section, we explain the states and weights notation used to develop the equilibrium statistical mechanical models used in this work. As a focus, we consider the case of an antibody binding to a single site on a receptor as shown in Fig S1A. We assume that the concentration of the antibody far exceeds that of the receptor, so that one binding event will not noticeably affect the concentration $c$ of free antibody.

The receptor can exist in two states where it is either unbound or bound to the antibody, where the relative probability of the bound state compared to the unbound state equals $\frac{c}{K_D}$, where $K_D$ is the dissociation constant of the receptor-antibody binding (1). The (normalized) probability of each state is given by its relative weight divided by the sum of the relative weights of all states. For example, the probability that the receptor is bound to the antibody is shown to be the standard $\frac{\frac{c}{K_D}}{1+\frac{c}{K_D}}$ sigmoidal response. As expected, the receptor will always be unbound in the absence of antibody ($c = 0$), while the receptor will always be bound at saturating concentrations of antibody ($c \to \infty$).

Each receptor state also has a relative activity (i.e. the activity of the receptor when it is in this state). In the context of EGFR, where the fractional activity is measured relative to the receptor in the absence of antibody, the relative activity of the unbound state is by definition equal to 1. As in the main text, we define the activity of the bound receptor to be $\alpha$, where $\alpha = 0$ implies that the receptor is completely inactive when the antibody is bound and $\alpha = 1$ represents the opposite limit where antibody binding does not inhibit the receptor's activity. A value of $0 < \alpha < 1$ represents an antibody that partially inhibits activity upon binding whereas an antibody with potency $1 < \alpha$ increases activity upon binding.

The activity of the receptor in each state is given by the product of its relative activity and the normalized probability of that state. Lastly, the average activity of the receptor is given by the sum of its activity in each state,

$$\text{Activity} = \frac{1 + \alpha \frac{c}{K_D}}{1 + \frac{c}{K_D}}. \tag{S1}$$

As a quick aside, we note that all the models considered in this work analyze binding reactions that quickly reach steady, and hence the equilibrium model derived above will should accurately describe such systems. Eq (S1) can also be derived by considering the dynamics of the system shown in the rates diagram Fig S1B. The unbound receptor (R) will switch to the bound state (R–Ab) at the rate $ck_{\text{on}}$ (recall that $c$ is the concentration of antibody) and subsequently unbind at a rate $k_{\text{off}}$. Hence, the system is governed by the two differential equations

$$\frac{d[\text{R}]}{dt} = k_{\text{off}}[\text{R–Ab}] - ck_{\text{on}}[\text{R}] \tag{S2}$$

$$\frac{d[\text{R–Ab}]}{dt} = ck_{\text{on}}[\text{R}] - k_{\text{off}}[\text{R–Ab}]. \tag{S3}$$

Since the total amount of receptor $[\text{R}_{\text{tot}}] = [\text{R}] + [\text{R–Ab}]$ is fixed, these two equations are equivalent, and either one can be solved to yield the dynamics of the system, namely,

$$[R] = c_1 e^{-(k_{\text{off}}+ck_{\text{on}})t} + [\text{R}_{\text{tot}}]\frac{k_{\text{off}}}{k_{\text{off}} + ck_{\text{on}}}. \tag{S4}$$

$c_1$ is fixed by the initial concentration of free receptor $[R]$ at $t = 0$, but regardless of its value, we see that the exponential term will shrink to zero at a time scale of $\frac{1}{k_{\text{off}}+ck_{\text{on}}}$, after which the system will be in steady state with $[R] = [\text{R}_{\text{tot}}]\frac{k_{\text{off}}}{k_{\text{off}}+ck_{\text{on}}}$. Defining the dissociation constant $K_D \equiv \frac{k_{\text{off}}}{k_{\text{on}}}$, the probability that any receptor will be unbound in steady state is given by

$$\frac{[R]}{[\text{R}_{\text{tot}}]} = \frac{1}{1 + \frac{c}{K_D}} \tag{S5}$$

**A**

| Receptor State | Relative Probability | Normalized Probability | Relative Activity | Weighted Activity |
|---|---|---|---|---|
| *All receptor states* | $Z_j$ | $\frac{1}{\sum_j Z_j}$ | $\alpha_j$ | $\frac{\alpha_j}{\sum_j Z_j}$ |
| 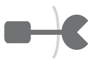 | $1$ | $\frac{1}{1+\frac{c}{K_D}}$ | $1$ | $\frac{1}{1+\frac{c}{K_D}}$ |
| 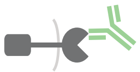 | $\frac{c}{K_D}$ | $\frac{\frac{c}{K_D}}{1+\frac{c}{K_D}}$ | $\alpha$ | $\alpha\frac{\frac{c}{K_D}}{1+\frac{c}{K_D}}$ |

**B**

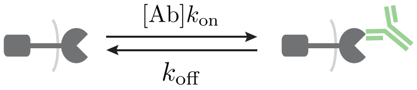


$$K_D \equiv \frac{k_{\text{off}}}{k_{\text{on}}}$$

**Figure S1. A statistical mechanical model of an antibody binding to a receptor.** (A) The relative probability and relative activity of each receptor state enable us to derive the average activity of the receptor. (B) This method gives the same result of the dynamics model shown by the multiple rates, provided that the system is in steady state.

as found in Fig S1A.

To close, we note that antibodies typically have a $K_D = 10^{-12}$–$10^{-8}$ M, and assuming a diffusion-limited on rate $k_{\text{on}} = 10^8$–$10^{-9}\,\frac{1}{\text{M}\cdot\text{s}}$, this implies that $k_{\text{off}} = 10^{-3}$–$10^1\,\frac{1}{\text{s}}$. Therefore, an upper bound (in the limit of little antibody) for the time scale it takes for such a system to reach steady state is $10^{-1}$–$10^3$ s, which is the time before experimental measurements should be conducted. Since antibodies are typically preincubated for even longer periods during experiments, an equilibrium model should be valid in all of the case studies we consider in this work.

#### A.2 The Fractional Activity of 3-Ab Mixtures

As described in the main text, the fractional activity of 2-Ab mixtures is given by Eq (2) if the two antibodies bind to distinct epitopes and Eq (3) if they bind to overlapping epitopes. These equations are straightforward to extend to mixtures with multiple antibodies by drawing all of the statistical weights and Boltzmann weights for the mixture (analogous to Fig 1) and then computing the fractional activity as per Section A.1.

For example, a mixture of three antibodies all binding to distinct epitopes would give rise to

$$\text{Fractional Activity}_{(1,2,3\text{ distinct epitopes})} = \left(\frac{1+\alpha_1\frac{c_1}{K_D^{(1)}}}{1+\frac{c_1}{K_D^{(1)}}}\right)\left(\frac{1+\alpha_2\frac{c_2}{K_D^{(2)}}}{1+\frac{c_2}{K_D^{(2)}}}\right)\left(\frac{1+\alpha_3\frac{c_3}{K_D^{(3)}}}{1+\frac{c_3}{K_D^{(3)}}}\right). \tag{S6}$$

If antibodies 1 and 2 bind to an overlapping epitope but antibody 3 binds to a distinct epitope, then

$$\text{Fractional Activity}_{(1,2\text{ overlapping epitopes; }3\text{ distinct epitope})} = \left(\frac{1+\alpha_1\frac{c_1}{K_D^{(1)}}+\alpha_2\frac{c_2}{K_D^{(2)}}}{1+\frac{c_1}{K_D^{(1)}}+\frac{c_2}{K_D^{(2)}}}\right)\left(\frac{1+\alpha_3\frac{c_3}{K_D^{(3)}}}{1+\frac{c_3}{K_D^{(3)}}}\right). \tag{S7}$$

If all three antibodies binds to overlapping epitopes, the fractional activity becomes

$$\text{Fractional Activity}_{(1,2\text{ overlapping epitopes; }3\text{ distinct epitope})} = \frac{1+\alpha_1\frac{c_1}{K_D^{(1)}}+\alpha_2\frac{c_2}{K_D^{(2)}}+\alpha_3\frac{c_3}{K_D^{(3)}}}{1+\frac{c_1}{K_D^{(1)}}+\frac{c_2}{K_D^{(2)}}+\frac{c_3}{K_D^{(3)}}}. \tag{S8}$$

#### A.3 Characterizing Ten EGFR Monoclonal Antibodies from Koefoed 2011

Koefoed *et al.* investigated how a panel of ten monoclonal antibodies inhibit EGFR by measuring the protein's activity at multiple antibody concentrations in the human cell line HN5 (2). By fitting these titration curves to Eq (1), we can infer the dissociation constant $K_D$ between each antibody and EGFR as well as the potency $\alpha$ of each antibody in the HN5 cell line (Fig S2A,B). For each curve, the $K_D$ corresponds to the midpoint of the curve (halfway between its minimum and maximum activity values) while $\alpha$ represents the activity at saturating antibody concentration.

Koefoed *et al.* found that the majority (7/10) of these antibodies reduced EGFR activity below 20% at saturating concentrations, and since mixtures of antibodies would likely further decrease activity their potency would be difficult to accurately measure. To that end, Koefoed *et al.* switched to the A431NS cell line that is partially resistant to EGFR antibodiess where they remeasured EGFR activity for all ten monoclonal antibodies as well as mixtures of two or three Abs (with 1:1 and 1:1:1 ratios, respectively). Each measurement was performed at a mixture concentration of $2\,\frac{\mu\text{g}}{\text{mL}}$, implying that each antibody was half as dilute in the 2-Ab mixtures and one-third as dilute in the 3-Ab mixtures relative to the monoclonal antibody measurement.

We assume that the antibody-EGFR binding interaction is identical in the A431NS cell line, and hence that these same $K_D$ parameters characterizes these antibodies in that cell line. Hence, by using

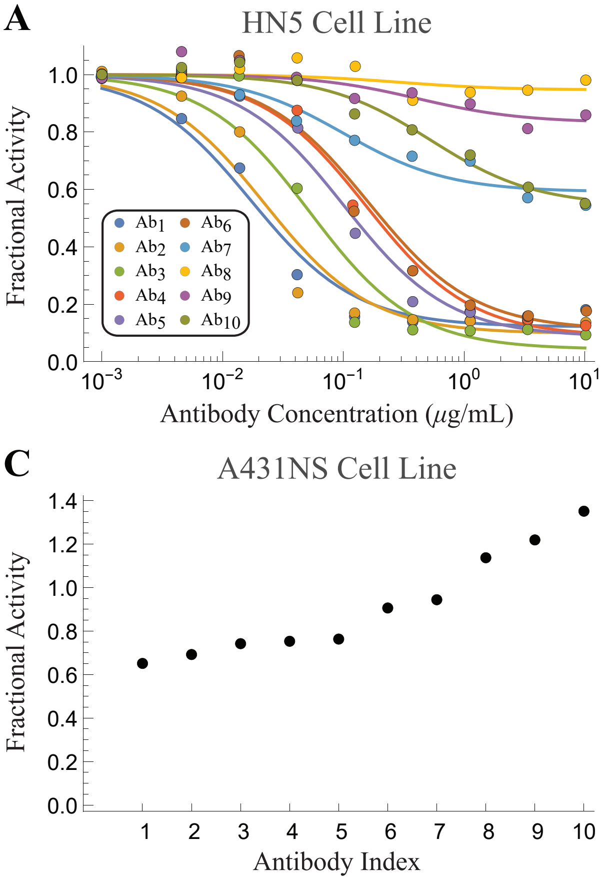


**B**

| Antibody Index | $K_D$ [$\mu$g/mL] (HN5/A431NS) | $\alpha$ (HN5) | $\alpha$ (A431NS) |
|---|---|---|---|
| $Ab_1$ | 0.02 | 0.12 | 0.65 |
| $Ab_2$ | 0.02 | 0.10 | 0.69 |
| $Ab_3$ | 0.05 | 0.04 | 0.74 |
| $Ab_4$ | 0.15 | 0.09 | 0.73 |
| $Ab_5$ | 0.10 | 0.09 | 0.75 |
| $Ab_6$ | 0.16 | 0.11 | 0.90 |
| $Ab_7$ | 0.10 | 0.59 | 0.94 |
| $Ab_8$ | 0.29 | 0.95 | 1.16 |
| $Ab_9$ | 0.41 | 0.83 | 1.26 |
| $Ab_{10}$ | 0.53 | 0.54 | 1.44 |

**Figure S2. Inferring the model parameters for the 10 monoclonal antibodies in Koefoed 2011.** (A) Activity of EGFR in the presence of ten antibodies in the HN5 cell line. (B) The inferred dissociation constant ($K_D$) and fractional activity in the presence of saturating antibody ($\alpha$) in each cell line. (C) Fractional EGFR activity in the presence of $2\,\frac{\mu\text{g}}{\text{mL}}$ of each antibody in the A431NS cell line. Data reproduced from Ref (2) Figure 2B,C.

the additional measurement of each antibody's potency at $2\,\frac{\mu g}{mL}$ in the A431NS cell line (Fig S2C), we can infer the potency $\alpha$ of each antibody in the A431NS cell line (Fig S2B). We propose no relationship between the $\alpha$ parameters in the HN5 and A431NS cell lines, and as noted by Koefoed *et al.*, some antibody mixtures may decrease activity in one cell line ($0 < \alpha < 1$) but increase activity ($1 < \alpha$) in the other cell line.

These parameters are sufficient to predict how any mixture (at any concentration and ratio) will behave in the HN5 and A431NS cell lines. Note that all data presented in the main text correspond to the A431NS cell line.

#### A.4 Comparing the HN5 and A431NS Cell Lines from Koefoed 2011

Koefoed *et al.* measured the potency of 176 mixtures in the A431NS cell line but only 55 mixtures in the HN5 cell line. Fig S3 extends our analysis to both the A431NS and HN5 cell lines. While the majority of mixtures have very little predicted and measured activity ($\lesssim 0.2$), approximately 13 outliers fall outside this range and appear to be poorly predicted.

While the coefficient of determination $R^2 = 0.61$ is significantly lower for this cell line, we note that: (1) there are far fewer data points in this cell line and (2) that our $R^2$ definition places more importance on points with larger predicted or measured fractional activity, and hence these few outliers have a disproportionate effect. That said, it remains unknown whether with more data our model would be as successful in the HN5 cell line. Another open question is why some antibody mixtures had inhibitory effects in one cell line but exacerbating effects in the other (e.g. the mixture of $Ab_3 + Ab_{10}$ resulted in 1.22 fractional activity of EGFR in the A431NS cell line but 0.19 fractional activity in the HN5 cell line).

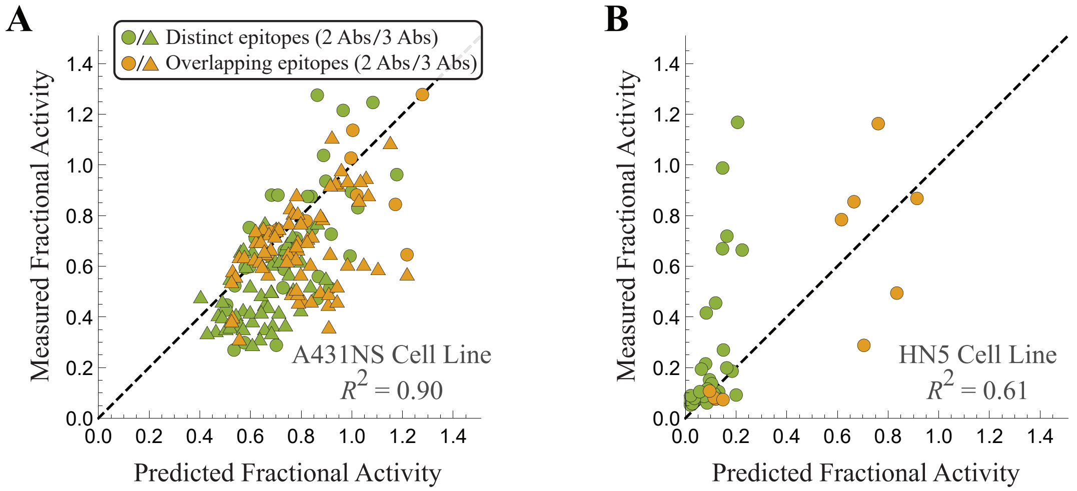


**Figure S3. Predicting the potency of antibody mixtures on EGFR in different cell lines.** Our model predictions versus experimental measurements for EGFR activity in the (A) A431NS and (B) HN5 cell lines.

#### A.5 Separating the 2-Ab and 3-Ab Predictions for EGFR Antibody Mixtures

Fig S4 separates the 2-Ab and 3-Ab mixture predictions from the three models in Fig 2. More specifically, Panels A and B of Fig S4 show the predictions for combinations of two and three antibodies using the epitope mappings produced by SPR (see captions on the diagonal of Fig 2A; there are four EGFR epitopes bound by antibodies #1-3, #4, #5-6, and #7-10, respectively).

Without this SPR data, we could have alternately assumed that antibodies all bind independently (Fig S4C,D) or that all antibodies vie for the same epitope (Fig S4E,F). Either of these models generate slightly worse predictions, as exhibited by their lower coefficients of determination $R^2$.

While the predictions for the A431NS cell line lie close to the diagonal, they are nevertheless systematically shifted to the right, indicating a larger predicted than measured fractional activity.

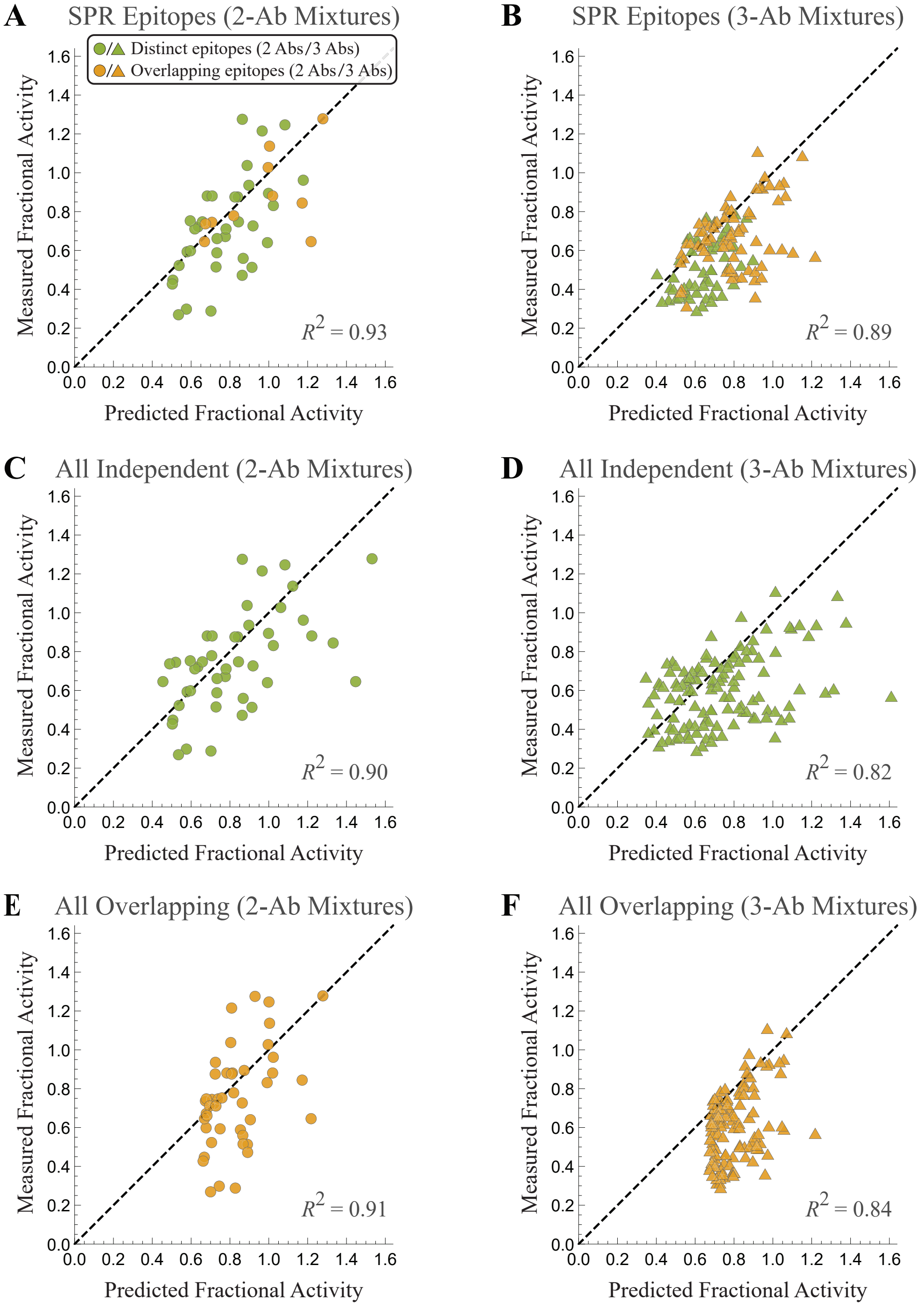


**Figure S4. Separate predictions for 2-Ab and 3-Ab mixtures.** (A,B) Predictions using the epitope mappings produced using SPR. (C,D) Predictions assuming that all antibodies bind independently. (E,F) Predictions assuming that all antibodies bind competitively.

To quantify this asymmetry, Fig S5 shows the distribution of measured minus predicted activity, demonstrating a tendency towards over-predicting activity.

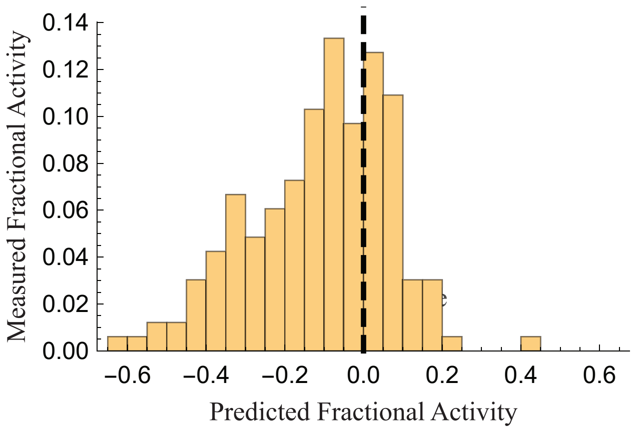


**Figure S5. Predicted minus measured fractional activity for all 2-Ab and 3-Ab mixtures.** The overall distribution is asymmetric about 0, showing a tendency to over-predict the fractional activity.

#### A.6 Comparing the 3-Ab Mixture Predictions using the different Epitope Mappings

In Fig 2B, we used the epitope mapping generated through SPR measurements to decide whether two Abs bound to distinct or overlapping epitopes. In contrast, Fig 3B shows the predictions when we instead inferred the epitope mapping from the 2-Ab activity data. We note that in this latter method, the 2-Ab mixtures were only used to group together antibodies that bound to the same epitopes (Fig 3A). Nevertheless, the cleanest test to determine how closely the two methods match one another is to compare their predictions for the activity of the 3-Ab mixtures which were not utilized in either case.

Fig S6 demonstrates that the two sets of predictions for the 3-Ab mixtures are nearly identical ($R^2 = 0.997$), demonstrating that there is essentially equal predictive power using either method to infer which antibodies bind to the same epitopes.

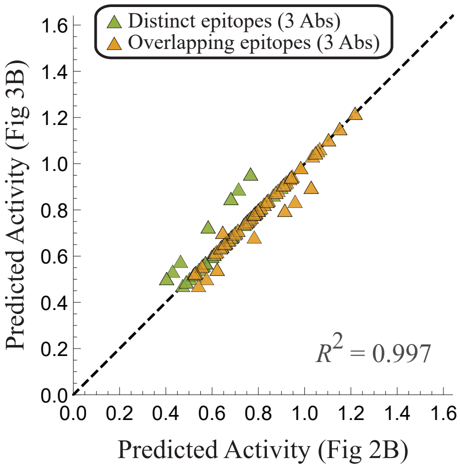


**Figure S6. Comparing the two sets of predictions for 3-Ab mixtures.** The predicted fractional activity of the 3-Ab mixtures from Fig 2B using the SPR epitope mapping ($x$-axis) are plotted against the predictions from Fig 3B using the 2-Ab activity data to infer the epitope map ($y$-axis).

#### A.7 A Continuum Model of Antibody Mixtures: Generalizing the Independent and Competitive Binding Models

Using the SPR measurements of the percent binding inhibition between each antibody pair (shown in Fig 4A), we defined the fraction $f = 1 - \frac{\%\text{ inhibition}}{100}$ of the doubly bound EGFR state for each antibody pair. For example, Abs #1 and #3 inhibit each other's binding by 100% ($f = 0$) whereas Abs #2 and #3 inhibit each other by only 90% ($f = 0.1$). This fraction $f$ multiplies the Boltzmann weight of the doubly bound state ($K_{D,\text{eff}}^{(2)} = \frac{K_D^{(2)}}{f}$ in Fig 1C), resulting in

$$\text{Fractional Activity} = \frac{1 + \alpha_1 \frac{c_1}{K_D^{(1)}} + \alpha_2 \frac{c_2}{K_D^{(2)}} + \alpha_1 \alpha_2 f_{12} \frac{c_1}{K_D^{(1)}} \frac{c_2}{K_D^{(2)}}}{1 + \frac{c_1}{K_D^{(1)}} + \frac{c_2}{K_D^{(2)}} + f_{12} \frac{c_1}{K_D^{(1)}} \frac{c_2}{K_D^{(2)}}} \tag{S9}$$

for a 2-Ab mixture. Without synergistic interactions, $f$ should lie between 0 and 1. Thus we assume that all values beyond this range are due to error and replace lower values with 0 and larger values with 1.

For 3-Ab mixtures, we assume that whenever any pair of antibodies (Abs $j$ and $k$) are simultaneously bound, they diminish the statistical weight of that state by $f_{jk}$. This yields the relation

$$\begin{aligned}&\text{Fractional Activity}\\ &= \frac{1 + \alpha_1 \frac{c_1}{K_D^{(1)}} + \alpha_2 \frac{c_2}{K_D^{(2)}} + \alpha_3 \frac{c_3}{K_D^{(3}} + \alpha_1\alpha_2 f_{12} \frac{c_1}{K_D^{(1)}} \frac{c_2}{K_D^{(2)}} + \alpha_2\alpha_3 f_{23} \frac{c_2}{K_D^{(2)}} \frac{c_3}{K_D^{(3)}} + \alpha_1\alpha_3 f_{13} \frac{c_1}{K_D^{(1)}} \frac{c_3}{K_D^{(3)}} + \alpha_1\alpha_2\alpha_3 f_{12} f_{23} f_{13} \frac{c_1}{K_D^{(1)}} \frac{c_2}{K_D^{(2)}} \frac{c_3}{K_D^{(3)}}}{1 + \frac{c_1}{K_D^{(1)}} + \frac{c_2}{K_D^{(2)}} + \frac{c_3}{K_D^{(3}} + f_{12} \frac{c_1}{K_D^{(1)}} \frac{c_2}{K_D^{(2)}} + f_{23} \frac{c_2}{K_D^{(2)}} \frac{c_3}{K_D^{(3)}} + f_{13} \frac{c_1}{K_D^{(1)}} \frac{c_3}{K_D^{(3)}} + f_{12} f_{23} f_{13} \frac{c_1}{K_D^{(1)}} \frac{c_2}{K_D^{(2)}} \frac{c_3}{K_D^{(3)}}}\end{aligned} \tag{S10}$$

where the triply bound state is maximally diminished by the multiplicative fractions $f_{12}f_{23}f_{13}$. Note that in the cases where all antibody pairs either bind purely competitively or independently ($f_{jk} \in \{0, 1\}$), Eq (S10) reduces to the appropriate form of fractional activity given by Eqs (S6)-(S8).

Like the purely independent/competitive model of antibody binding, the continuum model can predict the fractional activity of each antibody mixture *a priori* from the individual monoclonal antibody dose-response curves and the SPR blocking data. However, plotting the resulting model predictions (Fig S7A; $R^2 = 0.87$) resulted in poorer predictions than the original independent/competitive model (Fig 2B; $R^2 = 0.90$).

More precisely, mixtures containing only Abs #1-7 matched the model predictions far better (Fig S7B; $R^2 = 0.94$) than mixtures containing Abs #8, #9, or #10 (Fig S7C; $R^2 = 0.84$). Notably, Abs #8-10 were the only antibodies that individually increased activity (i.e. had potency $\alpha$ greater than 1 in the A431NS cell line as shown by Fig S2B), yet when they were combined with other antibodies they decreased EGFR activity. For example, while Abs #8 and #10 individually increase activity by 1.14 and 1.35, respectively, their mixture decreases activity to 0.65. This suggested that when Abs #8-10 are simultaneously bound with another antibody, the mechanism of action by which they increase activity may be disrupted. This idea is corroborated by the observation that antibody mixtures containing Abs #8-10 were systematically higher than the measured activity (Fig S7C).

To account for this behavior, we modified the continuum model so that when two antibodies are simultaneously bound (as in the final state in Fig 1C), their potencies are modified to the form $\alpha_{j,\text{eff}}$ (with one parameter per antibody, distinguished by the $j$ subscript). With this modification, Eq (S9) becomes

$$\text{Fractional Activity} = \frac{1 + \alpha_1 \frac{c_1}{K_D^{(1)}} + \alpha_2 \frac{c_2}{K_D^{(2)}} + \alpha_{1,\text{eff}} \alpha_{2,\text{eff}} f_{12} \frac{c_1}{K_D^{(1)}} \frac{c_2}{K_D^{(2)}}}{1 + \frac{c_1}{K_D^{(1)}} + \frac{c_2}{K_D^{(2)}} + f_{12} \frac{c_1}{K_D^{(1)}} \frac{c_2}{K_D^{(2)}}}. \tag{S11}$$

Similarly, all terms in Eq (S10) with products of two or more $\alpha_j$ become $\alpha_{j,\text{eff}}$.

Whereas the original continuum model required no fit parameters, the $\alpha_{j,\text{eff}}$ parameters must be inferred from the antibody mixture data. As a first step, we used the original potency parameters $\alpha_{j,\text{eff}} = \alpha_j$ for the first seven antibodies ($1 \le j \le 7$) and only fit the $\alpha_{j,\text{eff}}$ parameters for Abs #8-10.

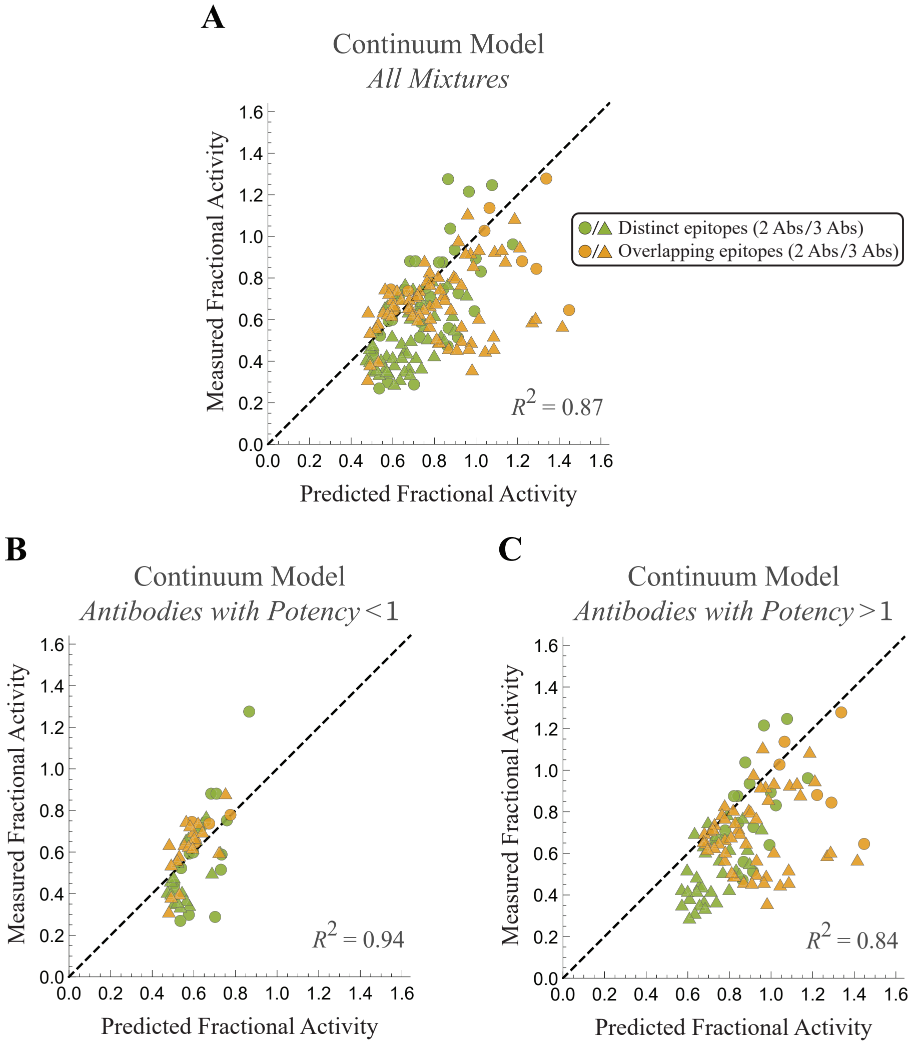


**Figure S7. The continuum model for antibodies with potency less than or greater than one.** (A) The predictions of the continuum model from Eqs (S9) and (S10) for all antibody mixtures. (B) Predictions for mixtures containing antibodies with potency less than one (i.e. mixtures composed only of Abs #1-7). (C) Predictions for mixtures containing at least one antibody with potency greater than one (i.e. mixtures must contain either Ab #8, #9, or #10 together with any other antibodies).

This led to a marked improvement in the model's ability to characterize the antibody mixtures (Fig S8B, $R^2 = 0.95$) compared to when no effective potency parameters were applied (Fig S8A, $R^2 = 0.87$). Altogether, this amounts to fitting a total of three parameters (one for each of the three antibodies with potency greater than 1; fit to the 109 antibody mixtures containing these three antibodies), with all other model parameters inferred directly from the data.

The model predictions can be further improved if an effective potency parameter $\alpha_{j,\text{eff}}$ is fit for all ten antibodies (Fig S8C, $R^2 = 0.98$), but only the effective potency of Abs #3 and #4 change by more than 15% from their individual potency values (Fig S8G), suggesting that the majority of these antibodies operate the same when individually bound to EGFR as when they are simultaneously bound with another antibody. This model has a total of ten parameters fit to the activity of all 165 antibody mixtures.

Lastly, we compared the continuum model against the original independent/competitive model by introducing the analogous $\alpha_{j,\text{eff}}$ parameters and characterizing each model's ability to characterize the activity of all antibody mixtures. While the continuum model with no effective potency parameters performed worse than the independent/competitive model (Fig S8D, $R^2 = 0.90$), the continuum model performed better when effective potency parameters were introduced for Abs #8-10 (Fig S8E, $R^2 = 0.94$) or for all ten antibodies (Fig S8F, $R^2 = 0.96$). Taken together, these results suggest that incorporating the spectrum of competition between antibodies can improve the ability to characterize antibody mixtures, and that antibodies (especially those that enhance EGFR activity) may behave differently when bound in combination with another antibody than when alone.

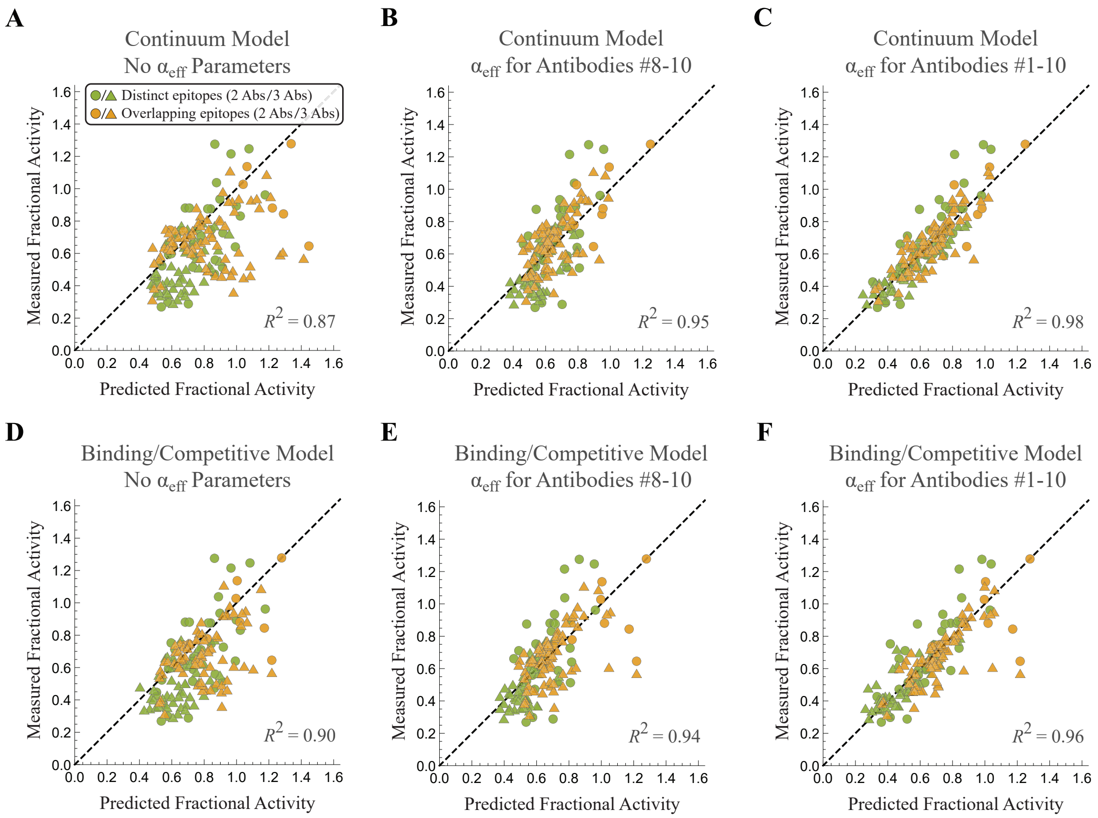


**G**

| Antibody Index | $\alpha$ (Panel A) | $\alpha$ (Panel B) | $\alpha$ (Panel C) | $\alpha$ (Panel D) | $\alpha$ (Panel E) | $\alpha$ (Panel F) |
|---|---|---|---|---|---|---|
| $Ab_1$ | 0.65 | 0.65 | 0.68 | 0.65 | 0.65 | 0.65 |
| $Ab_2$ | 0.69 | 0.69 | 0.76 | 0.69 | 0.69 | 0.70 |
| $Ab_3$ | 0.74 | 0.74 | 0.86 | 0.74 | 0.74 | 0.81 |
| $Ab_4$ | 0.73 | 0.73 | 0.36 | 0.73 | 0.73 | 0.43 |
| $Ab_5$ | 0.75 | 0.75 | 0.71 | 0.75 | 0.75 | 0.76 |
| $Ab_6$ | 0.90 | 0.90 | 1.00 | 0.90 | 0.90 | 0.98 |
| $Ab_7$ | 0.94 | 0.94 | 1.01 | 0.94 | 0.94 | 1.02 |
| $Ab_8$ | 1.16 | 0.60 | 0.60 | 1.16 | 0.60 | 0.62 |
| $Ab_9$ | 1.26 | 1.03 | 1.07 | 1.26 | 1.03 | 1.09 |
| $Ab_{10}$ | 1.44 | 0.97 | 0.95 | 1.44 | 1.02 | 1.06 |

**Figure S8. Comparing different modeling schemes for the 2-Ab and 3-Ab mixtures.** (A-C) The continuum model given by Eqs (S9) and (S10) where pairs of antibodies can partially inhibit one another's binding. (A) Using the potency parameters derived from monoclonal antibody dose-response curves (Fig S2B). (B) Using effective potency parameters $\alpha_{j,\text{eff}}$ for antibodies #8-10 when they simultaneously bind with another antibody. (C) Using effective potency parameters for all ten antibodies when they simultaneously bind with another antibody. (D-F) Analogous plots using the original binding model from Fig 2B where each antibody pair binds purely independently or competitively. (G) The resulting potency parameters used in each scheme.

#### A.8 Original Antibody Nomenclature

Table S1 shows the antibodies considered in this work were named in the original works of Refs. (2) and (3). In our work, we indexed the EGFR antibodies by their potency in the A431NS cell line and labeled the influenza antibodies by the viral group that they most effectively neutralized.

**Table S1. Matching the antibody nomenclature used in this work with the names in the original manuscript.**

| **Koefoed 2011 Antibody Nomenclature** | |
|---|---|
| *This Work* | *Original Work* |
| $\mathrm{Ab}_1$ | 1565 |
| $\mathrm{Ab}_2$ | 1320 |
| $\mathrm{Ab}_3$ | 1024 |
| $\mathrm{Ab}_4$ | 992 |
| $\mathrm{Ab}_5$ | 1277 |
| $\mathrm{Ab}_6$ | 1030 |
| $\mathrm{Ab}_7$ | 1284 |
| $\mathrm{Ab}_8$ | 1347 |
| $\mathrm{Ab}_9$ | 1260 |
| $\mathrm{Ab}_{10}$ | 1261 |

| **Laursen 2018 Antibody Nomenclature** | |
|---|---|
| *This Work* | *Original Work* |
| $\mathrm{Ab}_{\mathrm{A1}}$ | SD38 |
| $\mathrm{Ab}_{\mathrm{A2}}$ | SD36 |
| $\mathrm{Ab}_{\mathrm{B}}^{(1)}$ | SD83 |
| $\mathrm{Ab}_{\mathrm{B}}^{(2)}$ | SD84 |

### B Characterizing Distinct versus Overlapping EGFR Epitopes

In this section, we describe in more detail how we can use activity data from the 2-Ab mixtures to classify which subsets of antibodies bind to the same epitopes. A basic classification scheme would categorize two antibodies as binding to distinct epitopes if Eq (2) predicted their mixture's activity better than Eq (3); otherwise, the two antibodies would be categorized as binding to overlapping epitopes.

However, this simple classification scheme does not account for the uncertainty that arises from experimental noise. As an extreme case, if the activities of a mixtures are measured at extremely small concentrations, then the fractional activity will be $\approx 1$ for all mixtures (as will be the predictions from both the distinct and overlapping models), and this classification scheme would only be fitting the noise. Hence, it is best to measure the activity of each mixture at saturating antibody concentrations where the signal-to-noise of the system will be greatest.

We determined from Koefoed *et al.* (Figure S1) that the standard error of the mean (SEM) of their activity measurements was $\sigma = 0.04$, and we proceed to incorporate this uncertainty into our categorization scheme using a simple threshold model. More specifically, we add two components to the classification scheme: (1) As shown in Fig S9A, activity measurements that fall within $\sigma$ of the midpoint of the two model predictions are left unclassified, since experimental noise could easily lead to such points being incorrectly classified. (2) If the two model predictions lie sufficiently close (within $4\sigma$) to one another, then the uncertainty (from both the measurement and the model predictions) make it difficult to distinguish between the two models with certainty.

As mentioned above, for potent antibodies that are measured at saturating concentrations, the difference between the independent and overlapping binding models will be large, making it easier to definitively classify antibody epitopes. Koefoed *et al.* measured their 2-Ab combinations at a total concentration of $2\,\frac{\mu g}{mL}$ (and hence $1\,\frac{\mu g}{mL}$ for each antibody), and Fig S2A suggest that while this is close to saturating concentration for the majority of antibodies, increasing the concentration by 10x may have allowed more pairs of antibodies to be definitively categorized.

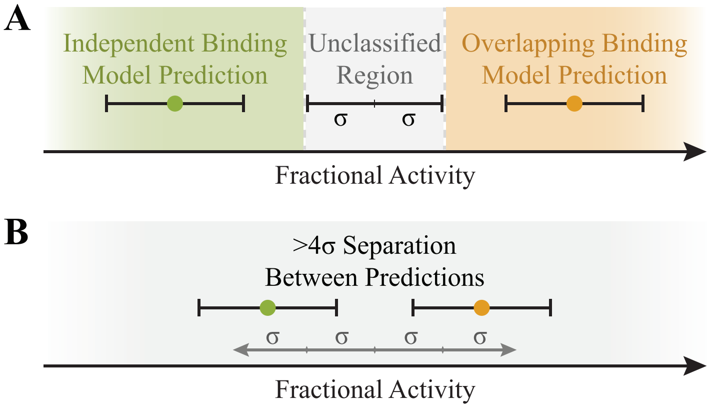


**Figure S9. Accounting for uncertainty in the classification scheme.** (A) 2-Ab combinations whose activity falls within $\sigma$ of the midpoint between the two model predictions will be labeled as "unclassified," since the difference in distance from the measurement to either prediction is less than the experimental error. (B) In addition, when the two model predictions are less than $4\sigma$ different, any measurement is labeled as "unclassified" because measurement error could result in overlap with both model predictions.

### C Characterizing Multidomain Antibodies

#### C.1 Relating Influenza Neutralization to Binding

In this section, we discuss how the microscopic dissociation constants of two tethered antibodies (shown in Fig 5C) relate to influenza viral neutralization. We begin by considering a single antibody with an effective dissociation constant $K_D^{(1)}$ that quantifies its avidity to the virus (4). The probability that this antibody at concentration $c$ will be bound to a virion is given by

$$p_{\text{bound}} = \frac{\frac{c}{K_D^{(1)}}}{1 + \frac{c}{K_D^{(1)}}}. \tag{S12}$$

For influenza virus with $N \approx 400$ hemagglutinin (HA) trimers, the number of bound trimers is given by $N_{\text{bound}} = N p_{\text{bound}}$.

The relationship between viral binding and neutralization remain unclear (5). It has been suggested that $\sim 50$ HA trimers are required to infect a cell (4). However, an IgG bound to one trimer may sterically preclude neighboring HA from binding. It has been proposed that neutralization is a sigmoidal function of neutralization (see Figure S1 of Ref (6)),

$$\text{Fraction Neutralized} = \frac{N^h + N_{50}^h}{N^h} \frac{\left(N p_{\text{bound}}\right)^h}{\left(N p_{\text{bound}}\right)^h + N_{50}^h}, \tag{S13}$$

where $N_{50}$ is the number of bound trimers required to reduce infectivity to 50%, $h$ is a Hill coefficient, and the prefactor assures that the fraction neutralized ranges from 0 (in the absence of antibody) to 1 (in the presence of saturating antibody).

In the absence of data for our influenza strain of interest, we will assume $h = 1$ in the following analysis. This enables us to rewrite Eq (S13) as

$$\text{Fraction Neutralized} = \frac{\frac{c}{\text{IC}_{50}^{(1)}}}{1 + \frac{c}{\text{IC}_{50}^{(1)}}}, \tag{S14}$$

where we have defined the inhibitory concentration of antibody at which 50% of the virus is neutralized

$$\text{IC}_{50}^{(1)} = \frac{N_{50}}{N + N_{50}} K_D^{(1)}. \tag{S15}$$

For example, if the virus is 50% neutralized when $N_{50} = 100$ trimers are bound, the midpoint of a viral neutralization curve would occur at roughly $1/5$ the antibody concentration required to bind 50% of the trimers, as has been observed for some influenza antibodies (see Figure 2 of Ref (7)).

We now consider the tethered two-domain antibody shown in Fig 5C. Denote the antibody concentration as $c$, the effective dissociation constants of its two domains as $K_D^{(1)}$ and $K_D^{(2)}$, and the effective concentration when both domains simultaneously bind as $\tilde{c}_{\text{eff}}$ (which we will shortly relate to the $c_{\text{eff}}$ in the antibody neutralization given in Eq (5)). The probability that this antibody is bound to a virion is given by

$$p_{\text{bound}} = \frac{\frac{c}{K_D^{(1)}} + \frac{c}{K_D^{(2)}} + \frac{c}{K_D^{(1)}} \frac{\tilde{c}_{\text{eff}}}{K_D^{(1)}}}{1 + \frac{c}{K_D^{(1)}}}. \tag{S16}$$

As above, we assume that neutralization is related to the binding probability through Eq (S13) with Hill coefficient $h = 1$, which upon substituting Eq (S16) yields

$$\text{Fraction Neutralized} = \frac{\frac{c}{\text{IC}_{50,\text{A1}}} + \frac{c}{\text{IC}_{50,\text{A2}}} + \frac{c}{\text{IC}_{50,\text{A1}}} \frac{c_{\text{eff}}}{\text{IC}_{50,\text{A2}}}}{1 + \frac{c}{\text{IC}_{50,\text{A1}}} + \frac{c}{\text{IC}_{50,\text{A2}}} + \frac{c}{\text{IC}_{50,\text{A1}}} \frac{c_{\text{eff}}}{\text{IC}_{50,\text{A2}}}} \tag{S17}$$

where we have defined the $\mathrm{IC}_{50}$s of both antibodies using Eq (S15) as well as the rescaled effective concentration

$$c_{\text{eff}} = \frac{N_{50}}{N + N_{50}} \tilde{c}_{\text{eff}}. \tag{S18}$$

Therefore, in the case where $h = 1$ the functional form of neutralization in Eq (S17) is identical to the probability that an HA trimer is bound given by Eq (S16), with the dissociation constants and the effective concentration rescaled by $\frac{N_{50}}{N+N_{50}}$.

To put these results into perspective, we note that many viruses are covered in spikes (analogous to influenza HA) that enable them to bind and fuse to their target cells (5), and hence the sigmoidal dependence between viral binding and neutralization is likely widely applicable. However, HIV is a clear exception, since each virion has an average of 14 envelope spikes (8). In that context, neutralization is roughly proportional to the number of bound spikes so that $\mathrm{IC}_{50} \approx K_D$ (9).

#### C.2 An Uninformed Null Model for Multidomain Influenza Antibodies

Fig S10 show the predicted versus measured $\mathrm{IC}_{50}$ for the influenza multidomain antibodies in Fig 6. To quantify how well this model does, we compared it to an uninformed null model that ignores the tethering component of the multidomain antibodies (i.e. the $c_{\text{eff}} \to 0$ limit). This null model treats all four antibody domains as binding to independent sites (although $\mathrm{Ab}_{\mathrm{A1}}$ and $\mathrm{Ab}_{\mathrm{A2}}$ have no affinity for the influenza B strains while $\mathrm{Ab}_{\mathrm{B}}^{(1)}$ and $\mathrm{Ab}_{\mathrm{B}}^{(2)}$ have no affinity to the influenza A1 or A2 strains). The results, shown in Fig S11, have larger predicted $\mathrm{IC}_{50}$ values since the avidity of the multiple domains is ignored, resulting in worse model predictions. In particular, the influenza B strains are very poorly characterized ($R^2 < 0$).

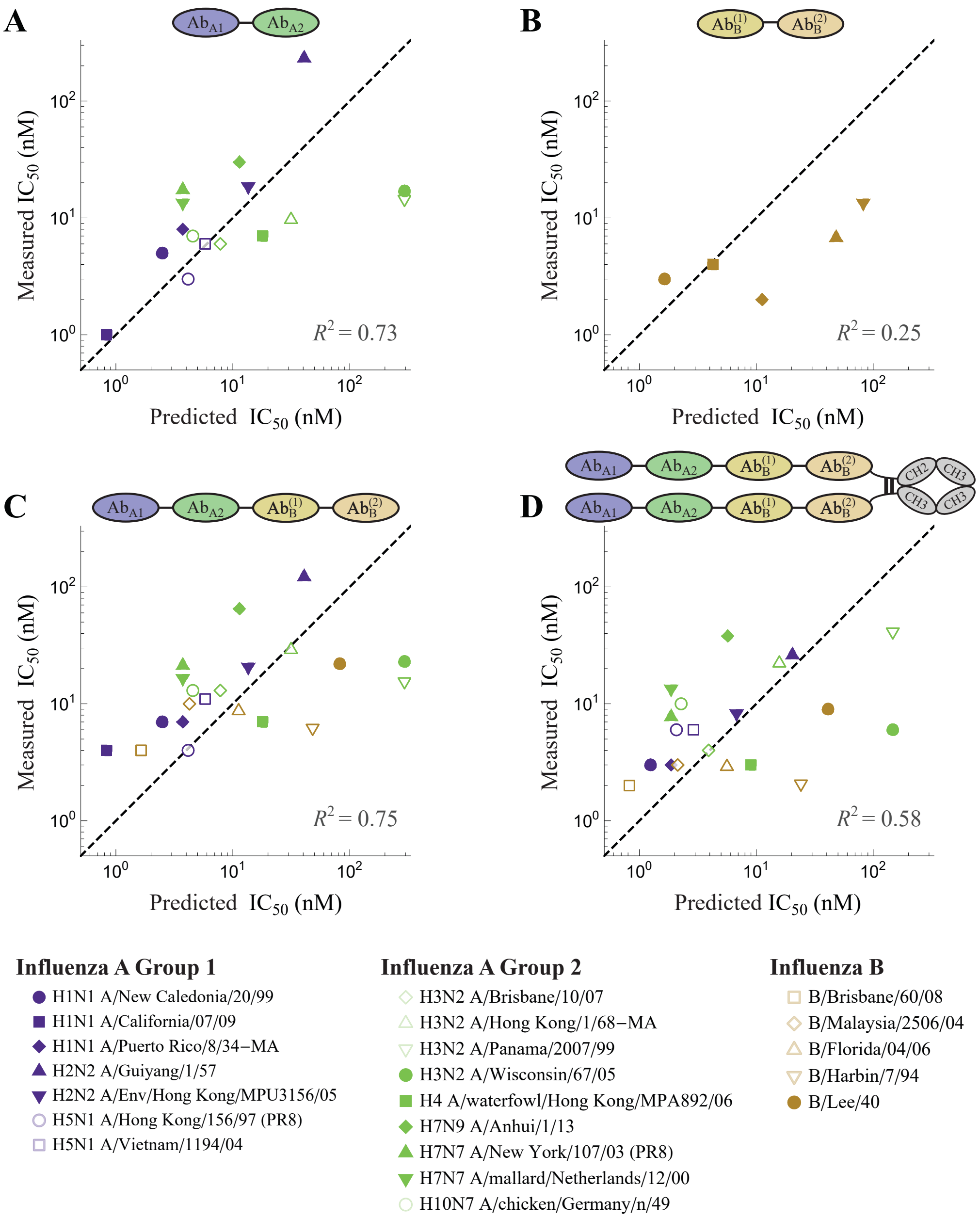


**Figure S10. The predicted versus measured $IC_{50}$ for the influenza multidomain antibodies in Fig 6.**

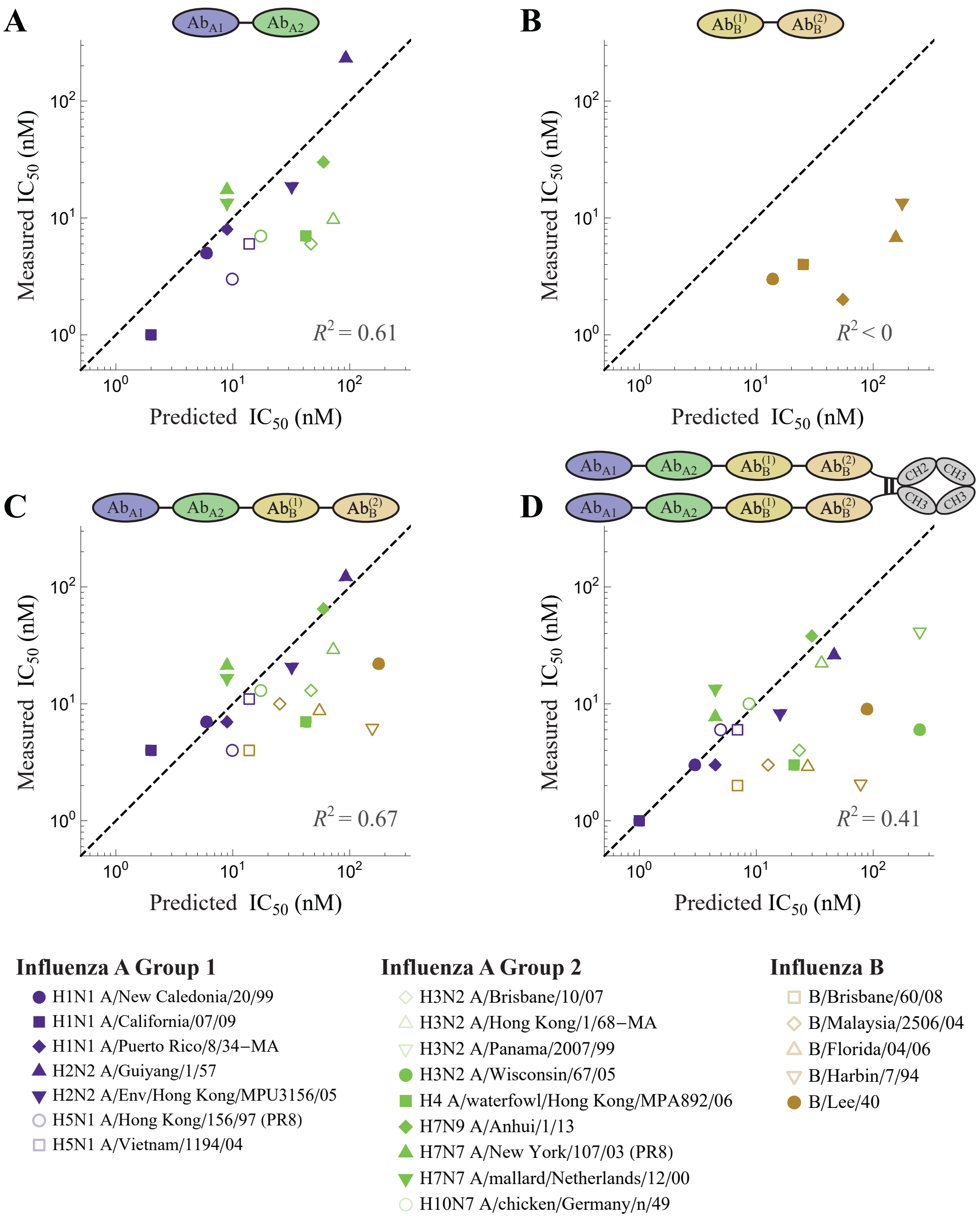


**Figure S11. A null model for multidomain influenza antibodies.** This model ignores the avidity between the different antibody domains and hence has larger $IC_{50}$ than Fig S10.

#### C.3 Assuming Different Antibody Constructs have Distinct Effective Concentrations

In the main text, we quantified the boost in avidity from tethering two antibodies using the effective concentration $c_{\text{eff}} = 1400\,\text{nM}$ by using least-squares regression to minimize the (log) predicted $IC_{50}$ for each tethered construct binding to all strains. This effective concentration depends on the distance between binding sites on a virion, and hence the tethered $Ab_{A1}$–$Ab_{A2}$ construct may have a different $c_{\text{eff}}$ when binding to influenza A group 1 and group 2 strains, and $Ab_B^{(1)}$–$Ab_B^{(2)}$ may have yet another effective concentration when binding to the influenza B strains.

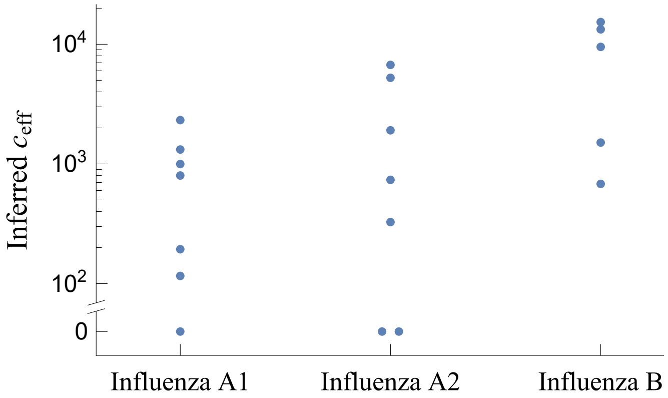


**Figure S12. Inferring $c_{\mathrm{eff}}$ between every tethered construct binding to each group of influenza virus.** The best-fit effective concentration of $\mathrm{Ab_{A1}}$–$\mathrm{Ab_{A2}}$ against the influenza A group 1 and group 2 strains, together with the effective concentration of $\mathrm{Ab_B^{(1)}}$–$\mathrm{Ab_B^{(2)}}$ binding to influenza B.

Fig S12 shows the best-fit $c_{\mathrm{eff}}$ for each of these cases. While this plot suggests that there are differences between each tethered antibody and influenza strain, we note that there is very limited data to infer such values (e.g., there are 7, 7, and 5 data points in the influenza A1, A2, and B groups, respectively; note that we ignore the H3N2 outlier strains A/Panama/2007/99 and A/Wisconsin/67/05 discussed in the main text). That said, incorporating this fine-grained level of modeling could further boost the accuracy of modeling efforts and is worth pursuing as more data is gathered.

Fig S13 shows the effect of characterizing the influenza multidomain antibodies using the geometric means of the non-zero $c_{\mathrm{eff}}$ values determined in Fig S12 for each viral strain. As expected, these yield better $\mathrm{IC_{50}}$ predictions, especially for the Influenza B strains.

Finally, we mention that Laursen *et al.* measured the efficacy of $\mathrm{Ab_{A1}}$–$\mathrm{Ab_{A2}}$ with different linkers of length 18 amino acids ($\sim$ 63Å), 38 amino acids ($\sim$ 133Å), and 60 amino acids ($\sim$ 210Å) against the four influenza strains H1N1 A/California/07/09, H1N1 A/Puerto Rico/8/34-MA, H5N1 A/Vietnam/1194/04, and H3N2 A/Wisconsin/67/05 (see Ref (3) Table S11). They found very little difference between the $\mathrm{IC_{50}}$ of each construct, which might naively suggest that the length of the linker does not matter. However, since these four strains were all negligibly inhibited by $\mathrm{Ab_{A2}}$ ($\mathrm{IC_{50}} \geq 1000\,\mathrm{nM}$; see Fig 6A), this domain cannot meaningfully contribute to bivalent binding, so that $\mathrm{Ab_{A1}}$–$\mathrm{Ab_{A2}}$ would be expected to be as potent as $\mathrm{Ab_{A1}}$ irrespective of linker length. On the other hand, the length of the linker should matter when both Abs in a multidomain antibody can bind, as is the case for $\mathrm{Ab_{A1}}$–$\mathrm{Ab_{A2}}$ binding to the three influenza A group 2 strains with $\mathrm{IC_{50}} < 1000\,\mathrm{nM}$ and for $\mathrm{Ab_B^{(1)}}$–$\mathrm{Ab_B^{(2)}}$ binding to the five influenza B strains.

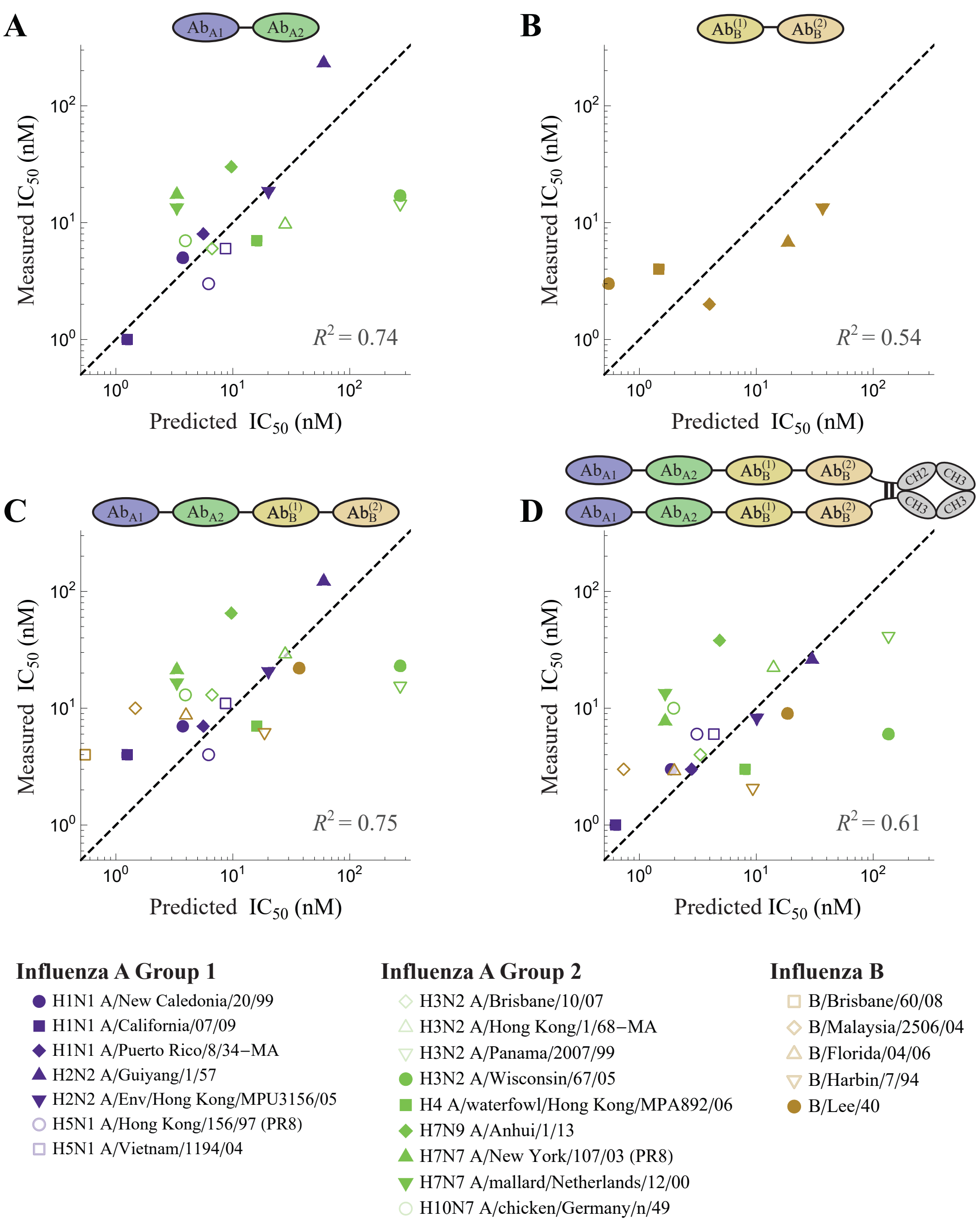


**Figure S13. Analyzing the influenza multidomain antibodies using a different $c_{\text{eff}}$ value per virus strain.** Parameter values were $c_{\text{eff}} = 600\,\text{nM}$ for the A1 strains, $c_{\text{eff}} = 1700\,\text{nM}$ for the A2 strains, and $c_{\text{eff}} = 4600\,\text{nM}$ for the B strains.

### References


1. Bintu, L., N. E. Buchler, H. G. Garcia, U. Gerland, T. Hwa, J. Kondev, and R. Phillips. 2005. Transcriptional regulation by the numbers: models. *Current Opinion in Genetics & Development.* 15:116–124.

2. Koefoed, K., L. Steinaa, J. N. Soderberg, I. Kjaer, H. J. Jacobsen, P.-J. Meijer, J. S. Haurum, A. Jensen, M. Kragh, P. S. Andersen, and M. W. Pedersen. 2011. Rational identification of an optimal antibody mixture for targeting the epidermal growth factor receptor. *mAbs.* 3:584–595.

3. Laursen, N. S., R. H. E. Friesen, X. Zhu, M. Jongeneelen, S. Blokland, J. Vermond, A. van Eijgen, C. Tang, H. van Diepen, G. Obmolova, M. van der Neut Kolfschoten, D. Zuijdgeest, R. Straetemans, R. M. B. Hoffman, T. Nieusma, J. Pallesen, H. L. Turner, S. M. Bernard, A. B. Ward, J. Luo, L. L. M. Poon, A. P. Tretiakova, J. M. Wilson, M. P. Limberis, R. Vogels, B. Brandenburg, J. A. Kolkman, and I. A. Wilson. 2018. Universal protection against influenza infection by a multidomain antibody to influenza hemagglutinin. *Science.* 362:598–602.

4. Xu, H., and D. E. Shaw. 2016. A Simple Model of Multivalent Adhesion and Its Application to Influenza Infection. *Biophysical Journal.* 110:218–233.

5. Klasse, P. J., and Q. J. Sattentau. 2002. Occupancy and Mechanism in Antibody-Mediated Neutralization of Animal Viruses. *Journal of General Virology.* 83:2091–2108.

6. Ndifon, W., N. S. Wingreen, and S. A. Levin. 2009. Differential Neutralization Efficiency of Hemagglutinin Epitopes, Antibody Interference, and the Design of Influenza Vaccines. *Proceedings of the National Academy of Sciences of the United States of America.* 106:8701–6.

7. Knossow, M., M. Gaudier, A. Douglas, B. Barrère, T. Bizebard, C. Barbey, B. Gigant, and J. Skehel. 2002. Mechanism of Neutralization of Influenza Virus Infectivity by Antibodies. *Virology.* 302:294–298.

8. Klein, J. S., and P. J. Bjorkman. 2010. Few and Far Between: How HIV May Be Evading Antibody Avidity. *PLoS Pathogens.* 6:e1000908.

9. Brandenberg, O. F., C. Magnus, P. Rusert, H. F. Günthard, R. R. Regoes, and A. Trkola. 2017. Predicting HIV-1 transmission and antibody neutralization efficacy in vivo from stoichiometric parameters. *PLoS Pathogens.* 13:1–35.
