## Supplementary material for "When two are better than one: Modeling the mechanisms of antibody mixtures": S2 File: ReadMe.txt

Supplementary Information for "When two are better than one: Modeling the mechanisms of antibody mixtures"

The supplementary Mathematica notebook "S1 File.nb" is organized into three parts:
(1) Program to Characterize Antibody Mixtures
 - This code can interact with the CSV files in this directory, enabling you to input your own antibody activity curves (or antibody parameters), categorize whether pairs of antibodies bind to distinct or overlapping epitopes, and predict the potency of an arbitrary mixtures
 - Each section in the notebook provides a detailed description on how to run the code. The CSV files provided in this same folder present a working example of how to input data, and these CSV files can be modified and the code rerun to analyze your own data
(2) Figures
 - Recreates all of the plots shown in this work
 - Note that the first time you evaluate any code (Shift+Enter) and are prompted to allow the Initialization cells to run, you must select "Yes" so that the antibody data can be properly loaded
(3) Initialization
 - Contains all of the data used in 
 - This section automatically runs the first time that you evaluate any piece of code in this notebook
